# NOVEL PRINCIPLES OF MOLECULAR GENETIC MAPPING OF THE INTERPHASE GENOME OF *Drosophila melanogaster*

**DOI:** 10.1101/2025.08.21.671599

**Authors:** Varvara A Khoroshko, Anton V Tsukanov, Nadezhda E Vorobyeva, Tatyana Yu Vatolina, Viktor G Levitsky, Igor F Zhimulev

**Affiliations:** Institute of Molecular and Cellular Biology, Siberian Branch of the Russian Academy of Sciences, Novosibirsk, Russia; Federal Research Center Institute of Cytology and Genetics, Siberian Branch of the Russian Academy of Sciences, Novosibirsk, Russia; Institute of Gene Biology, Russian Academy of Sciences, Moscow, Russia

## Abstract

Genome and chromosome maps have played a great role in the development of molecular genetics and biology. Gene activation, expression, and inactivation directly depend on the chromosomal (protein, nucleosomal) environment of the gene and formation of specific protein complexes on various gene structures. Polytene chromosomes are the only object in which interphase chromosomes can be analyzed, but the known *Drosophila* genome maps provide only an abstract view of gene distribution on a physical map, and there is no connection between these genes and the structures of interphase polytene chromosomes. A combination of bioinformatic methods was applied in this study; the data on interphase distribution of chromatin, H3K36me3 histone modifications, and the insulator protein Chriz, as well as the ChIP-seq, FAIRE-seq, and FISH methods, were used to investigate the genome-wide localization of key marker proteins. Having combined these mapping techniques for the small region 1AF of the X chromosome, we developed a novel method for analyzing the mutual arrangement of developmental and housekeeping genes, their promoters, and various types of proteins in the interphase genome and chromosome structures: compacted black bands, interbands, and gray bands, as well as sites of localization of exons and introns of housekeeping genes. Mapping was based on three consecutive stages: the 4 state Hidden Markov Model (hereinafter referred to as 4HMM) and data distribution for H3K36me3 and Chriz from the cells in which polytene chromosomes had been formed were used to localize interbands; FISH probes were then prepared from interband DNA, and blocks of developmental and housekeeping genes in interphase chromosome structures were mapped. The elaborated mapping methods can be further used to build similar maps for the entire interphase genome of *Drosophila*. Comparison of the map of bands and interbands in the region 1AF revealed full matching of the boundaries of black bands (developmental genes) and TADs. Within the regions where the housekeeping genes are located (the groups of interbands and gray bands), the TADs is formed on the basis of a gene cluster (the interband–gray band complex).

## Introduction

Interphase (polytene) chromosomes, in which the most active expression of genetic material takes place, are of particular interest for studying the organization of a functioning genome. During this very period, polytene chromosomes are in their most decondensed state and are accessible for being studied using molecular genetic methods and visual analysis.

A vast amount of data on localization of P-elements (Bourbon et al., 2002) and small deletions resulting from P-element transposition in chromosomes (Ryder et al., 2004) has been gained in *Drosophila* genetics. Essential functions have been identified in these studies (Peter et al., 2002). All these rearrangements can be very good markers, since their specific location on both physical and cytological maps is known. A polymorphic set of markers was used for human genome mapping: alternative forms of traits are supposed to exist for them, which can be described when performing analysis and constructing maps: for example, the known PDRF (restriction fragment length polymorphism) markers result from DNA mutations at restriction sites. Many human chromosome maps have been obtained by localization of genetic markers using the FISH method. Along with the known nucleotide sequences of the human genome, information is gained about the genetic markers such as probes and contigs, which are used to analyze genetic diseases. Bands in metaphase chromosomes, which arise as a result of differential staining performed using various techniques, are also mapped.

For gaining an insight into the genome sequence, it is important to both sequence DNA and construct physical maps of chromosomes with localization of chromosome structures: the ones densely packed (bands) and decondensed to various extents (interbands), genes, their structural and regulatory elements, as well as the chromatin state in them. For *Drosophila*, the map stored in the FlyBase database is characterized by significant inaccuracy and does not take into account the fact that the bands are subdivided into gray and black ones because the genes contained in them exhibit different activity levels. Moreover, the available maps do not show interbands, which are the crucial regions of the interphase genome where such processes as initiation, replication, and transcription occur (Zhimulev et al., 2014, 2024).

The maps of chromatin state distribution in the *Drosophila* genome are extremely important. Filion et al. used the DamID method for genome-wide mapping of 53 proteins in Kc167 *Drosophila* cell line (Filion et al., 2010). The group of proteins under study involved histone-modifying enzymes, nucleosome remodelers, components of general transcription mechanisms, insulator and heterochromatin proteins, structural components of chromatin, and DNA-binding factors. Five different chromatin states have been identified, which differed in terms of sets of proteins and histone modifications present in them. States corresponding to two types of active genes and chromatin inactivated by Polycomb group proteins or in heterochromatin have also been identified. Hence, the model contains two transcriptionally active states (RED and YELLOW), two inactive states (BLACK and BLUE), and heterochromatin (GREEN) (Filion et al., 2010).

For creating the next set of chromatin states, Kharchenko et al. carried out immunoprecipitation of 18 histone modifications and some proteins in S2 and BG3 cell lines. They identified a large number of states, and nine of those were of particular interest, each corresponding to different stages of transcription characterized by respective enrichment with a certain histone modification (Kharchenko et al., 2011).

Another chromatin classification is based on the general sensitivity of chromatin to DNase I (GCSDI), which identifies chromatin openness and accessibility for transcription (Milon et al., 2014). Using this method, Milon et al. analyzed the frequency of DNA breaks and determined the degree of chromatin openness for transcription in different genome structures (Milon et al., 2014).

In the aforementioned models, the authors focused on binding sites of proteins under study and different histone modifications without reference to the structures of polytene interphase chromosomes. In the 4HMM (Zhimulev et al., 2014, 2024; Khoroshko et al., 2016; Boldyreva et al., 2017) analysis of chromatin states involved three main types of the structures of interphase polytene chromosomes that are easily discernible using optical or electron microscopy and have been described a long time ago (Bridges, 1935, 1938). The black bands of densely packed transcriptionally inactive material correspond to the RUBY and MALACHITE chromatin state with a specific set of proteins and developmental genes. The RUBY and MALACHITE chromatin states reside within exons and introns of the developmental genes: in black bands of polytene chromosomes and in intergenic spacers. Each RUBY fragment is flanked by MALACHITE chromatin (Khoroshko et al., 2016, 2020). Open chromatin regions visualized by optical microscopy look more decondensed and light-colored. Those are the interbands persistently open for transcription (the AQUAMARINE state) hosting promoters of housekeeping genes and proteins of origin recognition complexes (ORCs), as well as gray bands (the LAZURITE state) hosting transcribed portions of gene bodies (exons and introns).

A total of 6562 housekeeping protein-coding genes and 3162 developmental genes have been identified in the *Drosophila* genome using 4HMM (FlyBase version 5) (Levitsky et al., 2020; Zhimulev et al., 2024). These groups of genes turned out to be independent within the same genome and differed at both the genetic and epigenetic levels. The following genetic difference (i.e., documented alterations in nucleotide sequences) is worth noting: promoters of both groups of genes have fundamentally different sets of motifs for transcription factor binding sites; in the genomes, they differ by size, lengths of introns and exons, intergenic spacers, the number of transcripts produced by a single gene, and the number of exons per gene. The epigenetic differences (i.e., those varying depending on the functional state of the genome) are related to replication timing, the degree of chromatin compaction and its elongation: H3K36me3 histone modification is responsible for regulation in housekeeping genes, while this mechanism is unknown for developmental genes (Zhimulev et al., 2014, 2023, 2024).

When constructing any map of the interphase organization of the eukaryotic genome, one needs to take into account the existence of three types of chromosome structures, chromatin states associated with them and two types of genes occupying these structures, promoters of housekeeping genes and their organization, as well as specific location of the sites where protein origin recognition complexes reside. The present paper addresses these questions.

Along with the data on gene location on the physical map, many molecular genetic methods that have never been used to study the genome organization in interphase chromosomes (in particular, the results of analyzing the 4HMM chromatin states and the ChIP-seq data on protein distribution as well as the FAIRE-seq data) were used for the first time in this paper. Having combined all these approaches, we developed the new stereotype of simultaneous high-precision mapping of bands and genes with respect to each other in the region 1AF, aiming to further perform whole-genome mapping.

## Materials and Methods

### Probe fabrication and fluorescence *in situ* hybridization (FISH)

The salivary glands of third-stage larvae were dissected using Ephrussi–Beadle modified saline, fixed in a 3:1 mixture of ethanol and acetic acid, squashed on a microscopic slide in 45% acetic acid, frozen in liquid nitrogen, and stored in 70% ethanol at –20°C. Fluorescence *in situ* hybridization (FISH) on polytene chromosomes was conducted according to the procedure described in ref. (Ashburner et al., 2005).

Twenty pairs of DNA probes were selected in accordance with the localization of 4HMM chromatin states corresponding to the interband regions and were obtained by PCR (Table 1). After being purified, the probes were labeled with Flu-12-dUTP and Tamra-5-dUTP (Biosan, Novosibirsk, Russia) in PCR with Klenow fragment (Vatolina et al., 2011). The chromosome specimens were analyzed by epifluorescence microscopy on an Olympus BX50 microscope; the images were recorded using a CCD Olympus DP50 digital camera system.

**Table 1.** Primers used for the study.

Chromatin immunoprecipitation (ChIP-seq) was performed and analyzed using exactly the same procedure as the one described previously (Vorobyeva et al., 2013; Mazina et al., 2018, 2021).

Dissection of *Drosophila* white prepupae (Oregon) was conducted manually in PBS/0.1% NP-40. The dissected salivary glands to be used for ChIP-Seq were collected into tubes containing PBS/0.1% NP-40 and placed onto ice. The dissected tissues were stored on ice for no longer than 1 hr. After the dissection, the tissues were heated to room temperature and fixed by adding FA to a concentration of 1% during 10 min. Fixation was stopped by adding 1/20V 2.5M Gly to the reaction during 5 min. After the fixation, a sonication buffer (ChIP lysis buffer) containing 0.1% SDS and PIC (Roche) was added to the tissues, and they were stored at –70°C. Ultrasonic treatment was conducted after thawing. The chromatin immunoprecipitation protocol was then performed as described previously (Vorobyeva et al., 2013). The precipitated DNA was used for library preparation using the NEBNext® Ultra™ II DNA Library Prep Kit for Illumina. The prepared library was sequenced on a NovaSeq6000 Sequencing System.

### The Hi-C data

The publicly available Hi-C datasets from the GEO database were used to analyze the spatial organization of chromatin in *Drosophila melanogaster*: GSM8188161 and GSM8188162 (Bing et al., 2024), GSM2679637 (Stadler et al., 2017), as well as GSM1863750 (Eagen et al., 2015). The dataset GSM2679637 corresponds to Stage 5 embryos, datasets GSM8188161 and GSM8188162 represent late embryonic Stage 14. GSM1863750 provides Hi-C interaction maps from polytene chromosomes, representing a highly specialized and rigid chromatin organization in differentiated salivary gland cells. Despite these biological differences, all datasets were processed using a unified computational pipeline to ensure consistency and comparability of the results.

### Hi-C processing

Raw paired-end FASTQ reads were cleaned and filtered using the fastp tool, version 0.24.0 (Chen, 2023). The following filters were utilized: (1) poly-G/poly-X trimming, where X is any nucleotide; (2) removing low-quality bases at both read ends (--cut_front, --cut_tail); (3) the minimal read length is 25 bp (--length_required 25); (4) quality control filtration with a threshold of 25 bp (--qualified_quality_phred 25); (5) quality correction based on overlapping (--correction) and automatic removal of adapters (--detect_adapter_for_pe).

The filtered reads were mapped to the masked genome of *Drosophila melanogaster* (dm6) using the BWA-MEM tool, version 0.7.18 (Li and Durbin, 2009); parameters: -SP5M -T0. The mapped reads were postprocessed using pairtools, version 1.1.2 (Open2C et al., 2023). It involved “pairtools parse” with alignment quality filtration (MAPQ ≥ 30), sorting (“pairtools sort”) and duplicate read removal (“pairtools dedup”). In order to increase the coverage depth and reduce intersample variability, the technical replicates were merged using “pairtools merge”, and duplicate reads were repeatedly removed.

The filtered read pairs were eventually utilized to obtain a contact matrix using the cooler tool, version 0.10.3 (Abdennur and Mirny, 2020). Only pairs with uniquely mapped reads were applied for creating the matrix. A matrix with a resolution of 1 kb was built (“cooler cload pairs”), and the matrices for other resolutions (2 kb, 4 kb, 8 kb, and 16 kb) were then calculated using “cooler zoomify”; all the matrices were normalized (balance parameter for “cooler zoomify”). TAD boundaries were determined according to the insulation score value (Crane et al., 2015), which was calculated using “cooltools” (Open2C et al., 2024) with the following parameters: “--min-frac-valid-pixels 0.75, -- min-dist-bad-bin 0, --ignore-diags 2 --threshold Li”. The insulation score calculation was performed for all resolutions, and in all cases the size of the sliding diamond-shaped window was determined as the product of the resolution and the coefficient (5, 10, 20, 40).

## RESULTS

As mentioned previously, according to the Bridges’ map, any chromosomal region can be partitioned into series of regions exhibiting shared characteristics: numerous large black bands interspersed with regions composed of groups of gray bands and interbands. The boundaries of these regions can be identified on the physical map by fluorescence *in situ* hybridization (FISH) using a DNA probe localized in the interband (the AQUAMARINE chromatin state) adjacent to the edge of a black band (the MALACHITE chromatin state marking the black band boundaries). For this reason, we have consistently been selecting FISH probes targeting interband DNA adjacent to black bands, such as probes CG3777-I and CG3777-II (Fig. 1C, F). The MALACHITE or RUBY chromatin states lying between the probes indicate that we have mapped the distal and proximal edges of a black band (Fig. 1); this finding was later confirmed by FISH. Conversely, if the LAZURITE chromatin state is detected between probes, then there are groups of gray bands and interbands (e.g., the region between probes CG3038 and CG3777-I (Fig. 1C, F). In cases where convenient FISH markers were unavailable, we used additional data, including the localization of housekeeping gene promoters (protein CHRIZ; Fig. 1D) or the histone modification H3K36me3 regulating transcriptional elongation of housekeeping genes (Fig. 1E).

**Figure 1.**
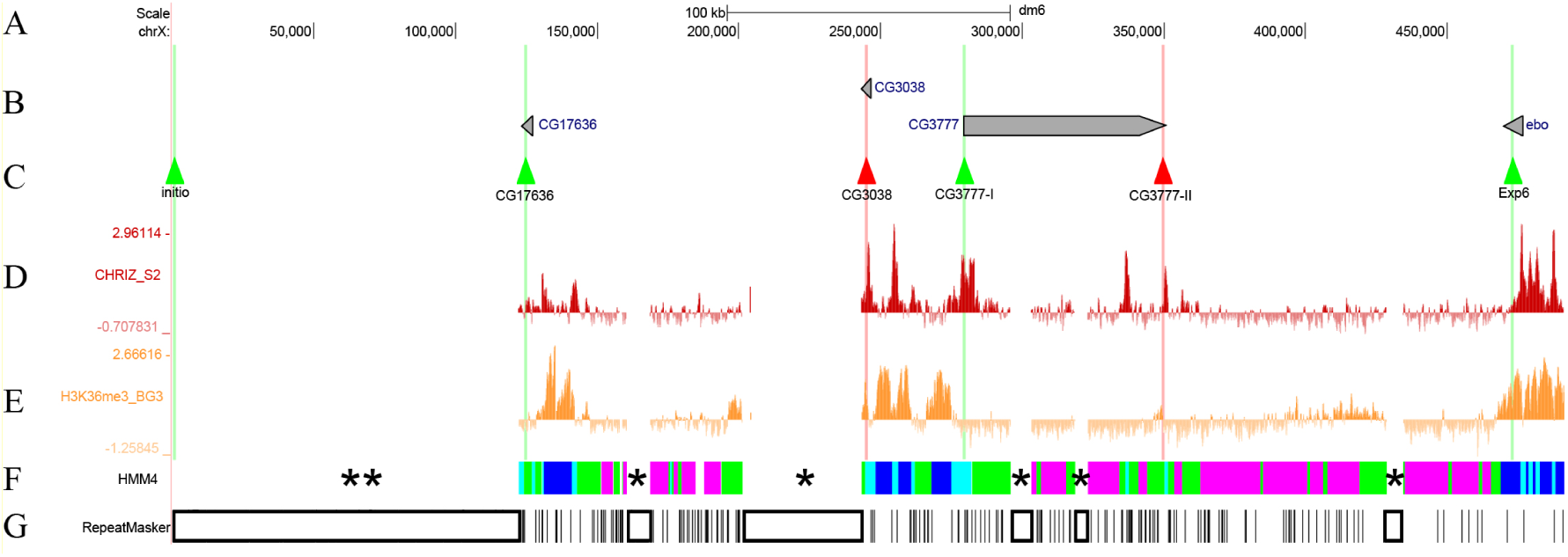
The scheme of selecting FISH probes in the region between chromosome start site 1A1 and 1B3-4. (A) The physical map and scale; digits denote the nucleotides; source: UCSC release dm6 (Haeussler et al., 2019). (B) Localization of genes carrying probes. (C) Localization of DNA probes, see details in the Materials and Methods section. (D) The distribution of interband-specific protein Chriz across S2 cells (Contrino et al., 2012). (E) The distribution of H3K36me3 histone modification specific for gray bands across BG3 cells (Contrino et al., 2012). (F) Localization of the four 4HMM chromatin states (Zhimulev et al., 2014); the asterisks denote data gaps corresponding to repeats according to the RepeatMasker program (see details below). (G) The data of the RepeatMasker program (Jurka, 2000) annotating repeats (localization of repeated sequences is shown with open triangles).

Gaps can sometimes be observed in the chromatin state map. They indicate the existence of different LINE, LTR, and satellite repeats, etc. (Fig. 1 F, G). Two asterisks denote the subtelomeric repeat; a single asterisk denotes repeat cluster regions whose precise sequencing is difficult to perform and is not presented in the releases on the physical map (Fig.1 F, G). There is a significant gap in all the data for band 1A5-6; it is caused by the presence of a 35 kb STR and the SAR2_DM repeat overlapping with it on the opposite strand. RepeatMasker is the program testing whether DNA sequences contain any repeats or low-complexity DNA regions. It yields a thorough annotation of repeats present in the requested sequence, as well as its modified version where all the annotated repeats are masked (Jurka, 2000). We can show using the RepeatMasker program that the data gaps in the 4HMM are actually DNA fragments that are rich in a large number of various repeats, which impedes chromatin state prediction using the model (Fig. 1 F, G). Taking these clarifications into account, we present the maps of chromosomal regions with sequence gaps and gaps in chromatin state maps (Fig. 1 F, G).

### Telomere

The telomeric/subtelomeric region of the X chromosome consists of open chromatin but does not exhibit a visible banding pattern and, in our view, is associated with the most distal thin band, 1A1 on the Bridges’ map) (Bridges, 1938). This region is approximately 122.2 kb long, and its material is poorly presented. It contains a large number of transpositions of retrotransposons belonging to three families: HeT-A, TART, and TAHRE; all of them refer to the class of LINE retroelements lacking long terminal repeats (Shpiz and Kalmykova, 2007; Kalmykova and Sokolova, 2023). This peri-telomeric material is underrepresented upon polytenization (Zhimulev, 1998), and molecular models cannot predict chromatin states (Fig. 1 F, G). Furthermore, it contains no genes. The telomere of the X chromosome actively binds the Chriz protein, which is specific for chromatin decompaction. Its recruitment to telomeric regions apparently leads only to the transition of closed chromatin to open chromatin (Pokholkova et al., 2018).

For localizing this region on the cytological map, we selected probes specific to the most distal part of the telomere (probe initio in Fig. 1) and to the sequence of the very first annotated gene (probe CG17636) (Fig. 2B, F-H; Fig. 3B; signals are shown with arrowheads). On the cytological map, these probes appear as two separate signals lying 123.4 kb apart, with gray loose material residing between them (Fig. 3A, B). Additionally, we labeled the HeT-A repeat and found that the label completely stained the chromosome region up to the very first black band 1A5-6 (Fig. 3B). The observed colocalization pattern manifests itself as changes in fluorescence color: yellow color appears as a result of superposition of the red signal from HeT-A and the green signals from DNA probes.

**Figure 2.**
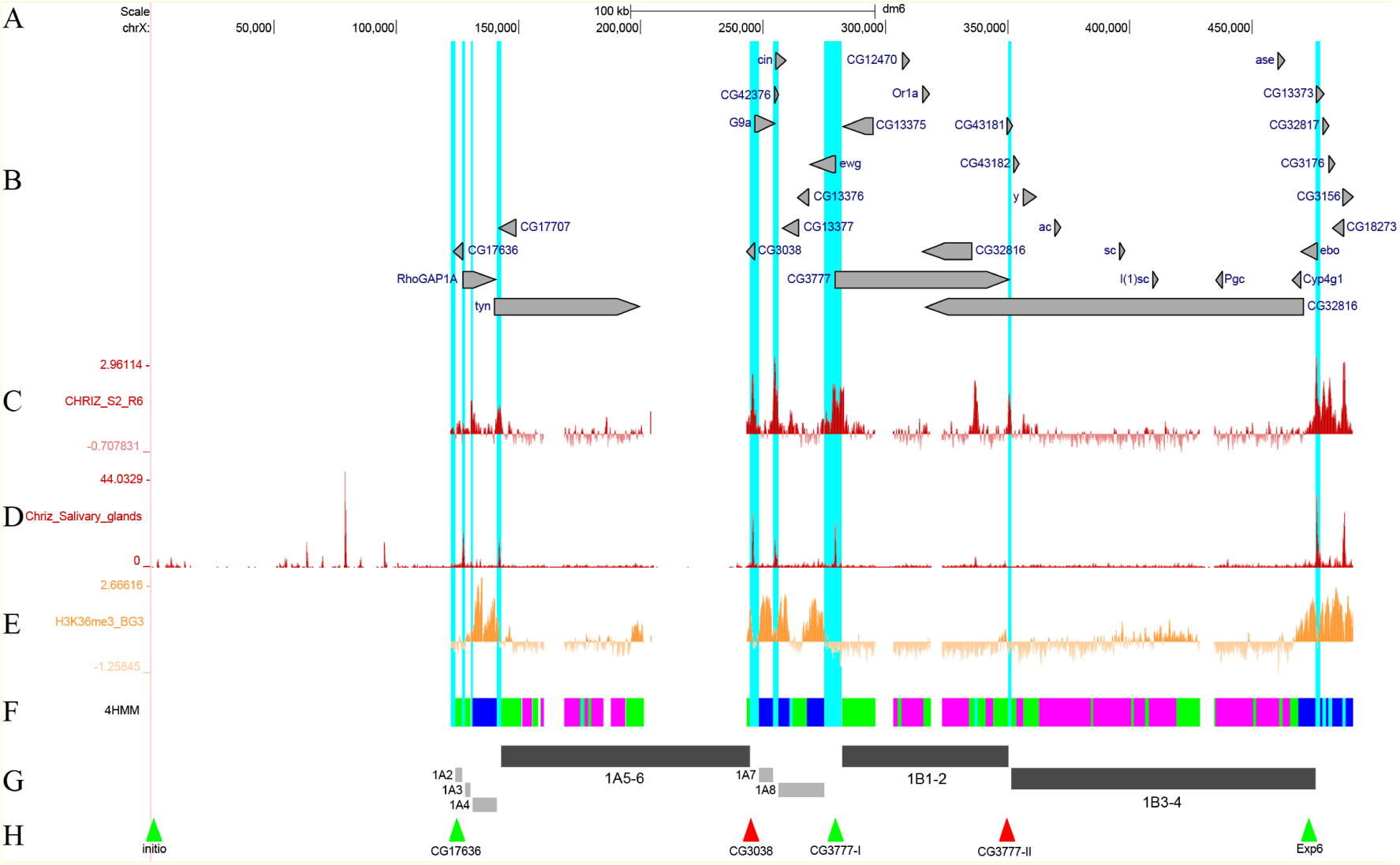
Mutual arrangement of the banding pattern of polytene chromosomes with genes and chromatin states in the region between the telomere and band 1B3-4. (A) Scale and molecular coordinates, UCSC release dm6 (Haeussler et al., 2019). (B) Localization and orientation of genes (the longest transcript is presented for each of them). (C) The distribution of interband-specific protein Chriz in S2 cells (Contrino et al., 2012). (D) The distribution of interband-specific protein Chriz in salivary gland cells. (E) The distribution of H3K36me3 in BG3 cells (Contrino et al., 2012). (F) The 4HMM of four chromatin states (Zhimulev et al., 2014). (G) Schematic localization of black and gray bands. (H) Localization of FISH probes. Light blue vertical lines show the interband regions for AQUAMARINE chromatin domains according to the 4HMM.

**Figure 3.**
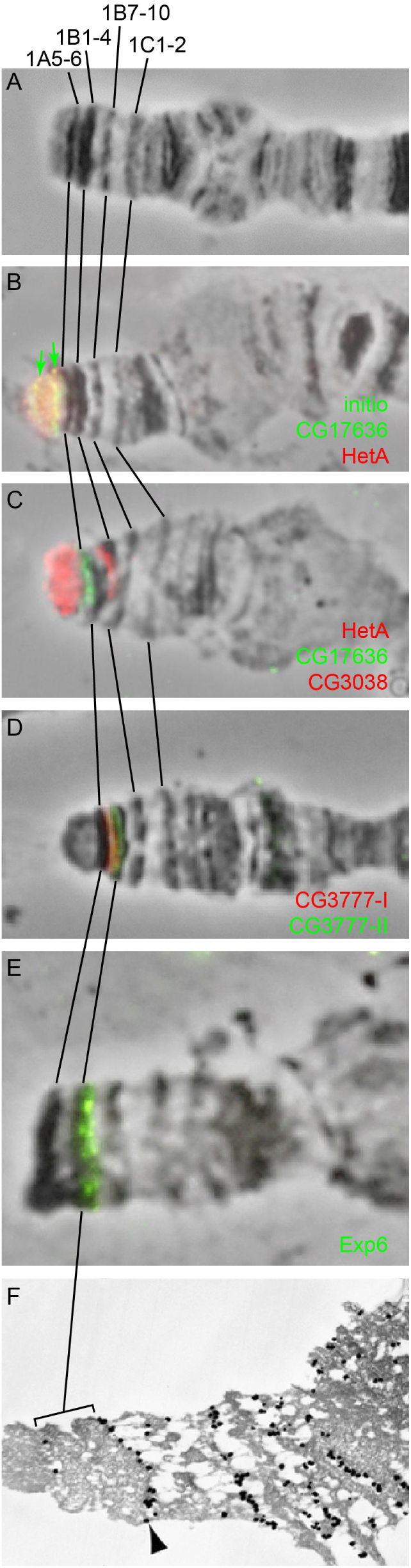
Cytological localization of FISH signals in the region between the telomere and band 1B3-4. (A) The map of the presented region. (B) Localization of probes initio, CG17636, and Het-A. (C) Localization of probes CG17636, CG3038, and Het-A. (D) Localization of probes CG3777-I and CG3777-II. (E) Localization of probe Exp6 (Demakova et al., 2020). (F) Electron microscopic localization of antibodies specific to gold-labeled MSL1 protein (the series of black dots shown with an arrowhead) (Semeshin et al., 2002).

### Mapping of black bands in the region 1AF

According to the electron microscopy data, thickness of individual bands in polytene chromosomes ranges from 0.3 μm to just below the limit of optical microscopy resolution (0.1–0.05 μm). Hence, based on the genome size, the lengths of the largest bands can be estimated to range from 100 kb to 5 kb for thin bands (Beermann, 1972). The more recent estimates of the lengths of large black bands are 167–190 kb; and 1.9–16.7 kb for thinner bands within the region between black bands 10A1-2 and 10B1-2 (Kozlova et al., 1994; Vatolina et al., 2011). Therefore, we focus on black bands and groups of gray bands and interbands separately.

### Region 1A2 - 1В3-4

For cytological mapping of the boundaries of black bands, we performed FISH of probes selected from 4HMM AQUAMARINE chromatin fragments, which bind antibodies against interband-specific protein Chriz in salivary gland cells of *Drosophila* and are adjacent to the material of black bands. These fragments are also characterized by high level of FAIRE-seq.

*Black band 1A5-6. A* single black band 1A5-6 immediately follows the telomeric region, from the sequencing initiation site toward the centromere, on the physical map (Fig. 2G, 3A). It is identified based on the massive accumulation of alternating RUBY and MALACHITE chromatin fragments, which are typical of black bands (Fig. 2F, G), being confined by the peak of Chriz protein distribution at the distal side. Length of the band, from the distal end (the boundary between AQUAMARINE/MALACHITE chromatin states in Fig. 2F) to the proximal boundary determined according to the coordinates of the CG3038 probe (Fig. 2H), is 101.5 kb according to the 4HMM. The distal boundary was denoted as the nearest AQUAMARINE with the 5’-end of the gene harboring Chriz in S2 cells. This mapping reveals more distally located three gray bands, which are consistent with the Bridges’ map well (the gray bands will be described below). The band 1A5-6 carries many aforementioned repeats (the gap in Fig. 2F). The 4HMM cannot predict chromatin states in the regions where repeats localize without data on nucleotide sequence; cytological localization is absent for proteins because of the large number of long repeats lying in this region (Fig. 1G). Therefore, the 4HHM model fails to predict chromatin states for the large ∼ 42 kb long fragment that resides within the band (chrX:201396-243368) and the smaller ∼ 8.4 kb long fragment (chrX: 160414-168815). The band contains the extensive *tyn* gene, and the shorter *CG17707* gene at the reverse strand (Fig. 3B).

*Black bands 1B1-2* and *1B3-4* are detected as two individual structures extremely rarely; they typically look as a single band (1B1-4 in Fig. 3A), although the Bridges’ map contains both of them: 1B1-2 and 1B3-4 (Bridges, 1938). Indeed, the conducted study revealed that these two bands contain alternating RUBY and MALACHITE chromatin fragments, which are typical of black bands, separated by an AQUAMARINE chromatin fragment where peaks belonging to the Chriz protein typical of interbands are also located (Fig. 2C, F). Furthermore, this region was also found to carry an insertion site of P-ICon (Demakov et al., 2010) based on the P-element, which is also inserted into open chromatin (Rubin and Spradling, 1982). Hence, the interband 1A8/1В1-2 has been successfully mapped. All this facts give grounds for considering it proved that there exists the tiny interband 1B1-2/1B3-4 (the length of the AQUAMARINE chromatin fragment is 1.2 kb) (Fig. 2). Small interbands are more likely to be detected in stretched chromosomes (this method was used by C. Bridges when constructing the map of the X chromosome (Bridges, 1938); therefore, it seems justified to identify the small interband and present it on the map in our study and on the C. Bridges’ map). For localizing the band, we used probes CG3777-I and CG3777-II that had been fabricated in an earlier study (Khoroshko et al., 2020) and resided in interbands 1A8/1B1-2 and 1B1-2/1B3-4 (Fig. 2F, G, Fig. 3D); they clearly determined the boundaries of band 1B1-2 and its length (67.9 kb) on the 4HMM physical map. The distal end of band 1B3-4 is determined by the proximal end of interband 1B1-2/1B3-4 (probe CG3777-II), while the proximal end, by the 3’ end of the *Exp6* gene (the *ebo* gene in dm6) (Fig. 2F-H), which is connected to band 1B3-4 chromatin (Fig. 2B, G; 3D). The *CG3777* gene entirely resides within band 1B1-2, while the remaining three genes are located within its intron, in the parallel and antiparallel strands. Figure 3F shows the results of electron microscopy mapping of antibodies specific to MSL1 protein, which bind to the transcribed regions of the genes of the X chromosomes (exons and introns or the gene body) (Kuroda et al., 2016). One can see that the signal from the probe runs along the boundary between band 1B3-4 and gray band 1B5, which contains the gene body and merges with band 1B3-4 (Demakova et al., 2020). The length of band 1B3-4 on the 4HMM physical map is 124.3 kb. The *CG32816* gene almost entirely lies within the 119.1 kb band 1B3-4. Eight other genes are also located in introns of this gene, residing entirely (*CG43182*, *y*, *ac*, *sc*, *l(1)sc*, *Pgcl*, *ase*, and *Cyp4g1*) on the parallel and antiparallel strands (Fig. 2B).

### Region 1B7-10…1D1-2

*Black band 1B7-10.* The black band 1B7-10 consists predominantly of RUBY and MALACHITE chromatin and carries the *Appl* and *vnd* genes; the *CG13366* gene at its right flank is linked to the band with its 3’ end, and its coding portion comprises a fragment of LAZURITE chromatin (Fig. 4B, F). Probes CG4293 and CG13366 were used for band localization; they are detected as two individual signals at the proximal and distal ends in interbands 1B6/1B7-10 and 1B7-10/1B9-10, respectively (Fig. 5B) and clearly define the boundaries of band 1B7-10 as well as its length (85.3 kb) on the 4HMM physical map.

**Figure 4.**
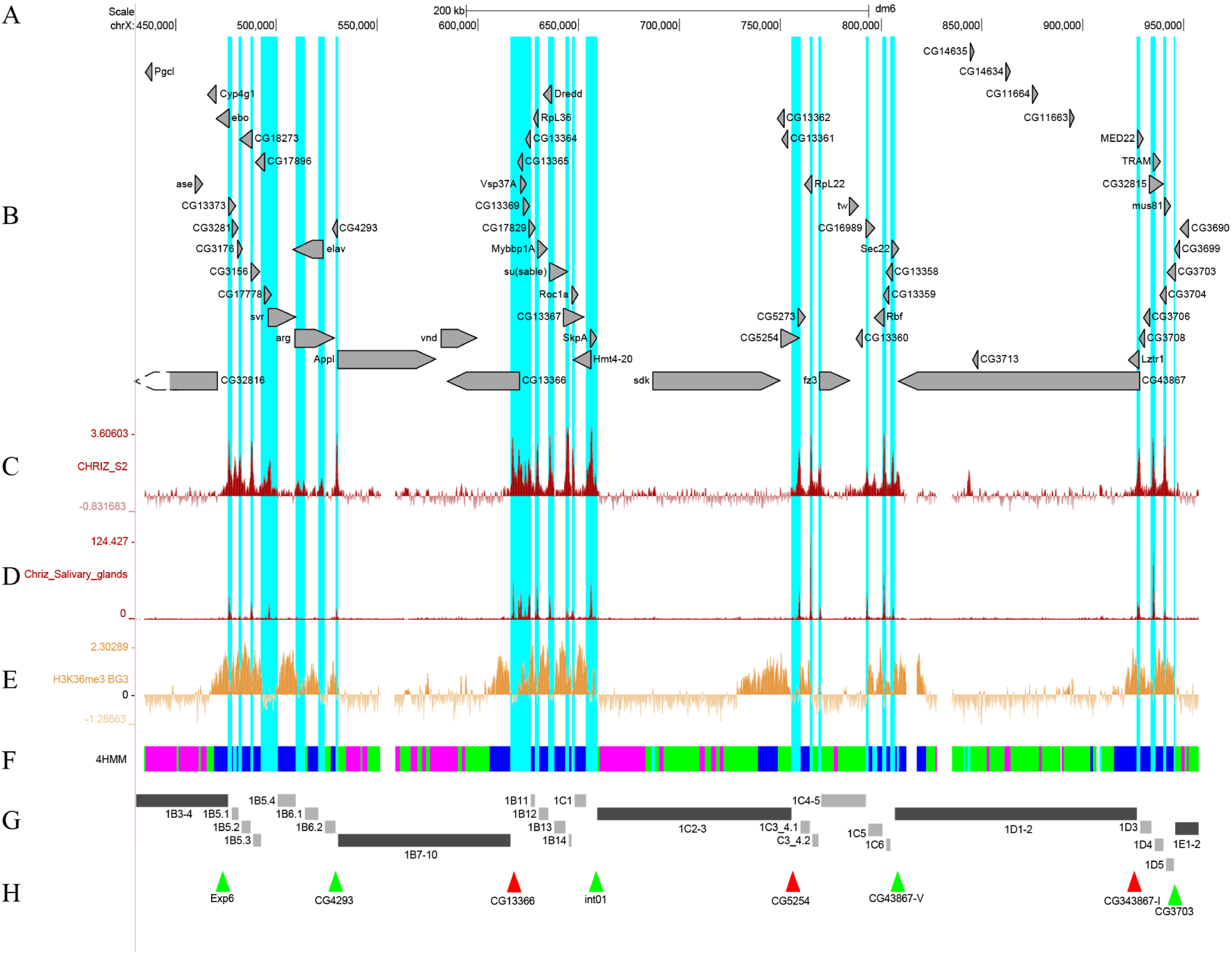
Mutual arrangement of the banding pattern of polytene chromosomes with genes and chromatin states within the region 1B7-10…1D1-2. (A) Scale and molecular coordinates, UCSC release dm6 (Haeussler et al., 2019). (B) Localization and orientation of genes (the longest transcript is presented for each of them). (C) The distribution of interband-specific protein Chriz in S2 cells (Contrino et al., 2012). (D) The distribution of interband-specific protein Chriz in salivary gland cells. (E) The distribution of H3K36me3 in BG3 cells (Contrino et al., 2012). (F) The 4HMM of four chromatin states (Zhimulev et al., 2014). (G) Schematic localization of black and gray bands. (H) Localization of FISH probes. Since there are no suitable genes for obtaining probes within the DNA region in the physical map more distal with respect to band 1C2-3, the probe was obtained from intergenic DNA and is referred to as int (intergenic). Light blue vertical lines show the interband regions for AQUAMARINE chromatin domains according to the 4HMM.

**Figure 5.**
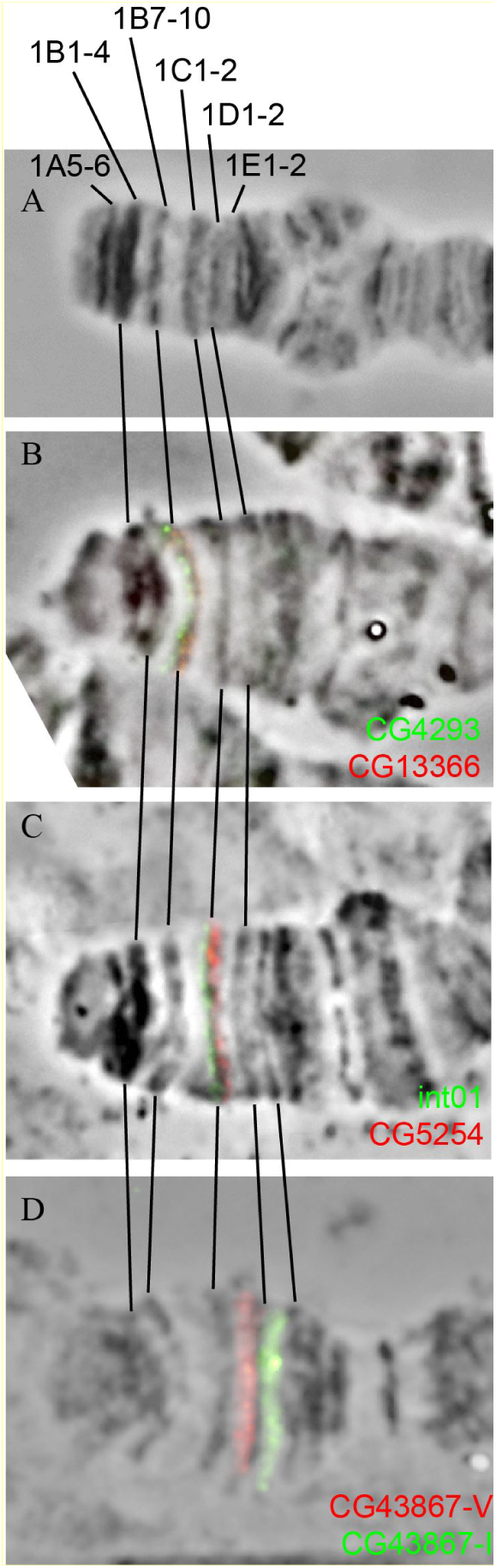
Cytological localization of FISH signals in the 1B7-10…1D1-2 region. (A) The map of the presented region. (B) Localization of probes CG4293 and CG13366. (C) Localization of probes int01 and CG5254. (D) Localization of probes CG43867-V and CG43867-I.

*Black band 1C2-3.* Band 1C2-3 consists of RUBY and MALACHITE chromatin (at an ∼ 1:1 ratio) and contains the *sdk*, *CG13362*, *CG5254*, and *CG13361* genes (Fig. 4B, F). The probes int01 and CG5254 were used for band localization; the probes are detected as two individual signals at the proximal and distal ends of the band in interbands 1C1/1C2-3 and 1C2-3/1C4-5, respectively (Fig. 4F-H, Fig. 5C), and clearly define the boundaries of band 1C2-3 and its length (96.1 kb) on the 4HMM physical map.

*Black band 1D1-2.* Band 1D1-2 consists predominantly of MALACHITE chromatin with small inclusions of RUBY chromatin, and carries the *CG43867*, *CG14634*, *CG1664*, and *CG11663* genes; the *Lztr1* gene at the right flank is linked to the band with its 3’ end, and its coding region comprises a fragment of LAZURITE chromatin (Fig. 4B, F-H). The second fragment of LAZURITE chromatin resides along the left edge of the band and is formed by the coding region of the *CG43867* gene (Fig. 4B, F). For band localization, we used probes CG43867-V and CG43867-I, which were obtained in a previous study (Khoroshko et al., 2020) and are detected as two individual signals within the proximal and distal ends of the band, respectively (Fig. 5D) and clearly define the boundaries of band 1D1-2 and its length (120.1 kb) on the 4HMM physical map.

### Region 1E1-2…2A1-2

*Black band 1E1-2.* The band 1E1-2 consists of RUBY and MALACHITE chromatin and carries the *CG3699*, *CG3690*, *CG18823*, *TfIIA-S-2*, *CG14631*, and *su(w[a])* genes as well as the 3’ ends of the *CG11638* and *CG32814* genes (Fig. 6 B, F). Probes CG3703 and CG3021 were used for band localization; they are detected as two individual signals at the proximal and distal ends of the band in interbands 1D4/1E1-2 and 1E1-2/1E3-4, respectively (Fig. 7C) and clearly define the boundaries of band 1E1-2 and its length (79.7 kb) on the 4HMM physical map.

**Figure 6.**
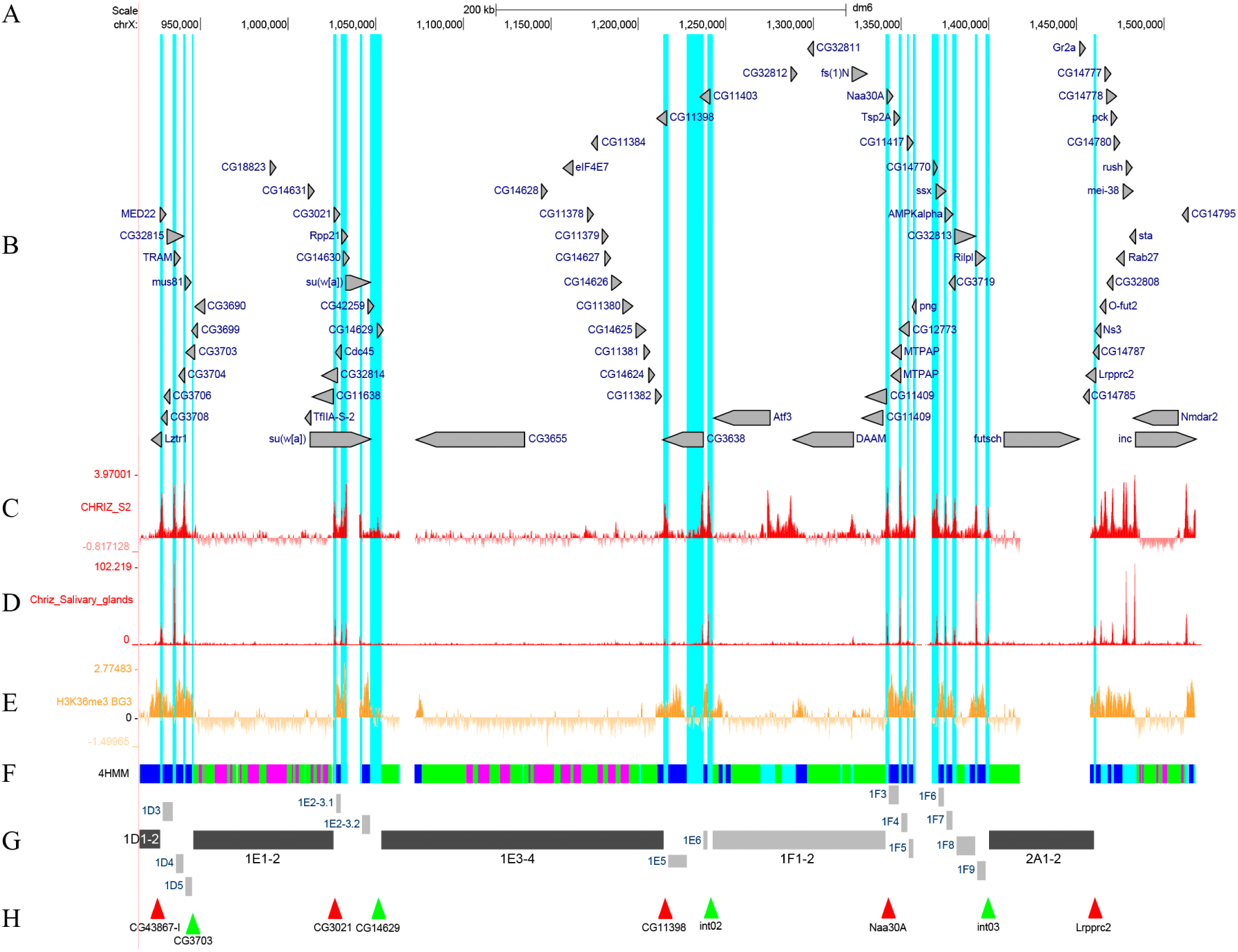
Mutual arrangement of the banding pattern of polytene chromosomes with genes and chromatin states within the region 1E1-2…2A1-2. (A) Scale and molecular coordinates, UCSC release dm6 (Haeussler et al., 2019). (B) Localization and orientation of genes (the longest transcript is presented for each of them). (C) The distribution of interband-specific protein Chriz in S2 cells (Contrino et al., 2012). (D) The distribution of interband-specific protein Chriz in salivary gland cells. (E) The distribution of H3K36me3 in BG3 cells (Contrino et al., 2012). (F) The 4HMM of four chromatin states (Zhimulev et al., 2014). (G) Schematic localization of black and gray bands. (H) Localization of FISH probes. Since there are no suitable genes for obtaining probes within the DNA region in the physical map more distal with respect to bands 1F1-2 and 2A1-2, the probe was obtained from intergenic DNA and is referred to as int (intergenic). Light blue vertical lines show the interband regions for AQUAMARINE chromatin domains according to the 4HMM.

**Figure 7.**
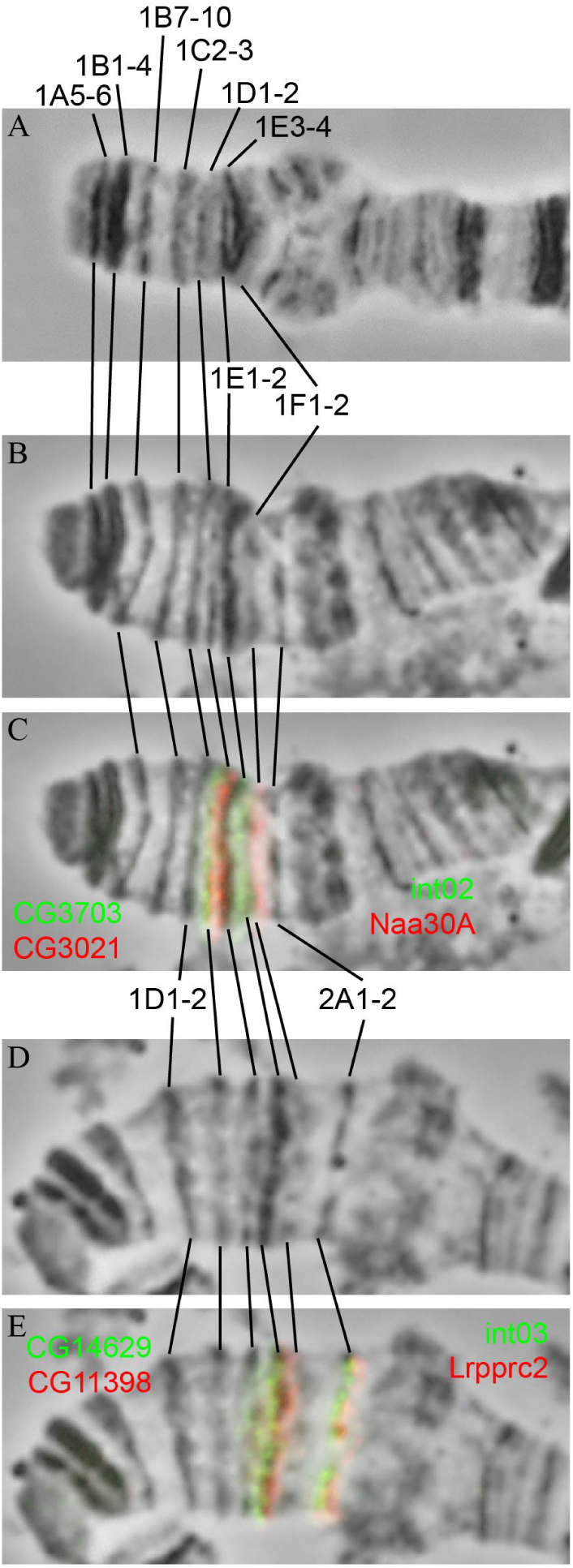
Cytological localization of FISH signals within the 1E1-2…2A1-2 region. (A, B) Localization of probes CG3703, CG3021, int02, and Naa30A. (C, D) Localization of probes CG14629, CG11398, int03, and Lrpprc2.

*Black band 1E3-4.* The band 1E3-4 consists of RUBY and MALACHITE chromatin and comprises 13 genes; the *CG11398* gene at the right flank is linked to the band via its 3’ end, and its coding region is a fragment of LAZURITE chromatin (Fig. 6B, F). Probes CG14629 and CG11398 were used for band localization; they are detected as two individual signals at the proximal and distal ends of the band in interbands 1E1-2/1E3-4 and 1E3-4/1E5, respectively (Fig. 7D) and clearly define the boundaries of band 1E3-4 as well as its length (161.1 kb) on the 4HMM physical map.

*The “black” band 1F1-2.* C. Bridges chose dense black bands when constructing the maps (Bridges, 1935, 1938) and for letter subdivisions; however, band 1F1-2 is neither dense nor black in this case. Taking into account the fact that this band opens letter subdivisions and has a large size (see below), we attributed it to the series of black bands. Band 1F1-2 consists of MALACHITE and LAZURITE chromatin and carries the *Atf3*, *CG32812*, *DAAM*, *CG32811*, *fs(1)N*, and *CG11409* genes (Fig. 6 B, F). The probes int02 and Naa30A were used for band localization; they are detected as two individual signals at the proximal and distal ends of the band in interbands 1E5/1F1-2 and 1F1-2/1F3, respectively (Fig. 7E) and clearly define the boundaries of band 1F1-2 as well as its length (98.5 kb) on the 4HMM physical map. In order to clearly delineate the region 1AF, we additionally localized the first band in the region 2AF. Band 2A1-2 contains the *futsch*, *Gr2a*, and *CG14785* genes; the *Lrpprc2* gene at the proximal end is linked to the band with its 3’ end, and its coding region comprises the fragment of LAZURITE chromatin (Fig. 6B, F). The 4HMM model displays the strongest data gap for this band (Fig. 1F, G) and a small MALACHITE chromatin fragment at its proximal end (Fig. 6 F, G). The probes int03 and Lrpprc2 were used for band localization; they are detected as two individual signals at the proximal and distal ends of the band in interbands 1F4/2A1-2 and 2A1-2/2A3-4, respectively (Fig. 7E) and clearly define the boundaries of band 2A1-2 as well as its length (59.7 kb) on the 4HMM physical map.

### Interband/gray band complexes

In earlier studies, the interbands were found to be no longer than 0.2 µm, while DNA length (bp) was considered negligible (Beermann, 1972). According to more recent findings, interband length is believed to vary from 0.6 to 3.0 kb in the domain between black bands 10A1-2 and 10 B1-2 (Vatolina et al., 2011) (see the data on thin bands above). Taking into account the findings, we describe bands and interbands within this domain.

In the gaps between densely compacted black bands, there lies a decompacted band and interband material. The black band material is compacted 160-to 220-fold; gray band material, 20-to 75-fold; and interbands, 5-to 10-fold [Kozlova et al., 1994; Vatolina et al., 2011]. The housekeeping genes occupy two cytological structures. Whereas the interband contains protein complexes of the origins of replication and the promoter of these genes, as well as binds the interband-specific insulator protein Chriz, the thin band lying downstream contains bodies (exons and introns) of these genes and binds the modified H3K36me3 histone, which is specifically detected in the transcription elongation sites. Gray bands are devoid of Chriz protein; anti-RNApolII antibodies are detected in transcribed exons and introns of the housekeeping genes. Therefore, the interband/gray band (AQUAMARINE and LAZURITE chromatin in 4HMM map) structure looks as a single functional unit (Zhimulev et al., 2014, 2024).

Figure 8 shows the data on localization of markers of the material open for replication and transcription (interbands and gray bands of polytene chromosomes) within the region under study.

**Figure 8.**
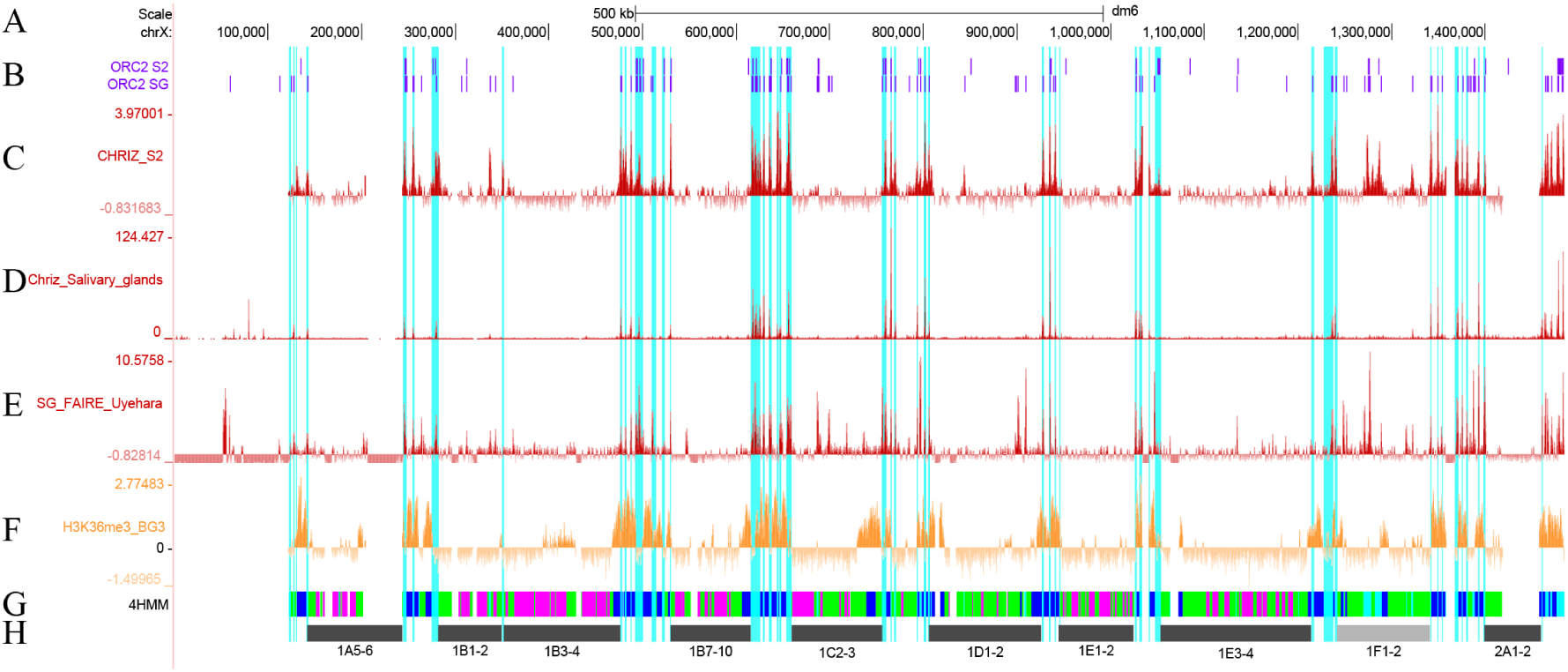
Detection of interband / gray band complexes within the 1A2…2A1-2 region. Scale and the molecular coordinates, UCSC release dm6 (Haeussler et al., 2019). (B) The distribution of protein complexes from the regions of Origin Recognition Complexes (ORC2) in S2 and salivary gland cells (Nordman et al., 2011). (C) The distribution of interband-specific protein Chriz across S2 cells (Contrino et al., 2012). (D) The distribution of interband-specific protein Chriz across salivary gland cells (the present study). (E) The distribution of open chromatin according to the FAIRE-seq procedure (Uyehara et al., 2022). (F) The distribution of H3K36me3 histone modification across BG3 cells (Contrino et al., 2012). (G) The 4HMM four chromatin state model (Zhimulev et al., 2014). (H) The localization coordinates of black bands within the region. Light blue vertical lines show the interband regions for AQUAMARINE chromatin domains according to the 4HMM model.

We compared the locations of interband-specific Chriz protein in cell culture and in polytene chromosomes of salivary glands, where the banding pattern was studied simultaneously for the entire mapped region 1AF. It is clear that the peaks of detected signals of both types virtually coincide. They also coincide with that for 4HMM AQUAMARINE chromatin (Fig. 8).

One can see that between the precisely mapped edges of black bands (Fig. 8H), there reside groups of interbands and gray bands (shown with light blue vertical lines in the figure). The positions of all the protein complexes being compared are well colocalized, which does not mean that there is 100% matching. The regions of maximally open chromatin (interbands) labeled with Chriz protein obtained for the *Drosophila* salivary gland cells (Fig. 8D) typically contain open chromatin at the sites of localization of FAIRE-seq fragments (Fig. 8E), as well as 4HMM AQUAMARINE and LAZURITE chromatin (Fig. 8G), and protein origin of replication complexes ORC2 obtained for the same cells (Fig. 8B). Modified H3K36me3 histone molecules are somewhat shifted with respect to the light blue lines (Fig. 8F), which is not surprising, since it labels gene bodies and corresponds to fragments of the LAZURITE chromatin state (Fig. 8G).

Hence, between the black bands, there reside series of interbands and gray bands formed by various portions of genes located in them. A total of nine such series have been detected: between the telomere and band 1A5-6; between bands 1A5-6 and 1B1-2; between bands 1B3-4 and 1B7-10; between bands 1B7-10 and 1C2-3; between bands 1C2-3 and 1D1-2; between bands 1D1-2 and 1E1-2; between bands 1E1-2 and 1E3-4; between bands 1E3-4 and 1F1-2; and between bands 1F1-2 and 2A1-2.

First and foremost, focus should be placed on some technical aspects related to mapping of thin loose bands. On the Bridges’ map (Bridges, 1938), gray bands are shown to be very thin and are often invisible even by electron microscopy. The model experiments, when the lengths of bands emerging from transposons with a known composition and size of constituent DNA were determined, demonstrated that DNA transposons ∼ 20 kb long and longer most typically yield bands easily visible by electron microscopy. DNA transposons ∼ 10 kb long and shorter most often do not form bands discernible by electron microscopy (Semeshin et al., 1989). Further discussion will demonstrate that the size of at least some gray bands is comparable to the microscopy resolution threshold. Therefore, 4HMM was used when the length of gray band was comparable to microscopy resolution: the AQUAMARINE/LAZURITE chromatin complexes corresponding to the interband/gray band pair were identified. With allowance for these considerations, we studied the gray bands within the region.

### Interband/gray band complexes between the telomere and band 1A5-6

The interval between the terminal end of telomeric repeat and the distal boundary of black band 1A5-6 is 20.6 kb long. According to the Bridges’ map, this region contains three gray bands 1A2, 1A3, and 1A4 (Bridges, 1938), each corresponding to the length of 6.8 kb on average. Their lengths lie at the threshold of electron microscopy resolution. The physical map shows that three genes reside in this region (*CG17636*, *RhoGAP1A*, and *tyn*, Fig. 2B), with three light blue vertical lines marking the position of Chriz protein. The most distal line, which is connected to the 4HMM AQUAMARINE chromatin fragment, belongs to the telomere (see above) and is not linked to the genes (Fig. 2B). The next two light blue lines toward the centromere contain promoters of the *CG17636* and *RhoGAP1A* genes, which are read in opposite directions (Fig. 2B). The H3K36me3 regions lie proximally and distally with respect to these lines, being indicative of transcription of the bodies of bidirectional genes (Fig. 2E). Hence, among the three bands under consideration (1A2, 1A3, and 1A4), the third band is the thinnest and is noticeably smaller than the other ones; we cannot see it. The other two interband/gray band gene complexes are on average 10.3 kb long. The final band resides upstream of band 1A5-6 and is associated with the promoter region of the *tyn* gene.

### Between bands 1A5-6 and 1B1-2

The interval between the terminal ends of bands 1A5-6 and 1В1-2 is 37.8 kb long. According to the Bridges’ map, it contains two gray bands 1A7 and 1A8 (Bridges, 1938); i.e., each interband/gray band complex has a length of 18.9 kb. These lengths are close to the generally accepted band length (30 kb) and are supposed to be easily detectable (Beermann, 1972; Sorsa, 1988a, 1988b).

On the physical map, this region contains eight genes (*CG3038*, *G9a*, *CG42376*, *cin*, *CG13377*, *CG13376*, *ewg*, and *CG3777*); six of them are connected to interbands. Three light blue lines pass through this region, correlating with Chriz protein in S2 and salivary gland cells, FAIRE-seq, and the 4HMM AQUAMARINE chromatin state; promoters of the genes (*CG3038*, *G9a*, *CG42376*, *cin*, *ewg*, and *CG3777*) lie within the interbands, and the gene bodies reside in the adjacent bands (Fig. 2).

Hence, on average, two interband/gray band complexes contain six genes residing entirely within them and two genes whose promoters are located within interbands, but their bodies reside in the black bands. Good agreement with H3K36me3 was found for the gene bodies (*G9a*, *CG42376*, *cin*, *CG13377*, *CG13376*, *ewg*, and *CG3777*), being indicative of their transcription. Hence, this region contains two gray bands and three interbands, all of them being visible; the region morphology corresponds to the Bridges’ map.

### Between bands 1B1-2/1B3-4

As mentioned earlier, the interband between black bands 1B1-2 and 1B3-4 is rarely detected, so both black bands look as a single band. Nevertheless, extremely rarely and mostly on stretched chromosomes (the method being used by C. Bridges when constructing revised maps), it is discernible and is present on the map; a 1.2 kb fragment of AQUAMARINE chromatin (Fig. 2) can be seen on the molecular map, with the Chriz peak localized in S2 cells, being virtually absent in salivary gland cells and FAIRE-seq (Fig. 8).

### Between bands 1B3-4 and 1B7-10

The interval between the terminal ends of bands 1B3-4 and 1B72 is 54.6 kb long; on the Bridges’ map, it contains two gray bands, 1B5 and 1B6 (Bridges, 1938), each of them supposedly corresponding to a length of 27.3 kb on average. These lengths are close to the generally accepted band length of 30 kb (see above). In this region, seven AQUAMARINE chromatin fragments can be identified on the physical map, containing interband-specific proteins Chriz and FAIRE-seq (Figs. 4 and 8). This fact can indicate that six gray bands are present. Each of the light blue lines corresponds to the promoter region of the gene; the gene bodies correspond to the H3K36me3 regions; and for each of the genes, there is a corresponding pair of 4HMM AQUAMARINE/LAZURITE chromatin states. Therefore, we assume that seven pairs of interband/gray band complexes containing the promoter and the gene body can be present. We have notionally referred to them as 1B5.1, 1B5.2, 1B5.3, 1B6.1, 1B6.2, and 1B6.3.

### Between bands 1B7-10 and 1C2-3

The interval between the terminal ends of black bands 1B7-10 and 1C2-3 is 43.4 kb long and contains five gray bands 1B11, 1B12, 1B13, 1B14, and 1C1 on the Bridges’ map (Bridges, 1938), each of them corresponding to a length of 8.6 kb on average. On the molecular map, within this region, one can see six fragments of AQUAMARINE chromatin with well-defined Chriz and FAIRE-seq peaks, and 5’ ends of active genes located in them, indicating that interbands are present (Fig. 4). Six Chriz and FAIRE-seq peaks are colocalized with AQUAMARINE chromatin; fragments of LAZURITE chromatin filled with H3K36me3 reside between them. Hence, all five gray bands can be precisely localized.

### Between bands 1C2-3 and 1D1-2

The interval between the terminal ends of black bands 1C2-3 and 1D1-2 is 51.2 kb long and contains a single gray band 1C4-5 on the Bridges’ map (Bridges, 1938); however, a detailed analysis of the molecular map of this region demonstrates that there are five or six pairs, ranging from the smallest interband/gray band pairs to the largest ones (light blue vertical lines in Figs. 4, 8). Therefore, it should be considered that this region contains bands of different lengths, some of them are not presented as gray bands on the cytological map. If 22.2 kb corresponding to band 1C4-5 is removed from the occupied intercept, the remaining gray bands are supposed to correspond to ∼ 7.2 kb, which will be clearly below the optical microscopy resolution. On the molecular map, one can see well-defined Chriz peaks in this region and the 5’ ends of active genes localized in them, indicating that interbands are present (Fig. 6). Six Chriz and FAIRE-seq peaks are colocalized with AQUAMARINE chromatin; between them, there lie LAZURITE chromatin fragments filled with H3K36me3 (Figs. 4, 8).

### Between bands 1D1-2 and 1E1-2

The interval between the terminal ends of black bands 1D1-2 and 1E1-2 is 19.2 kb long and contains two gray bands 1D3 and 1D4 on the Bridges’ map (Bridges, 1938), each of them corresponding to a length of 9.6 kb on average. On the molecular map, this region contains three light blue lines where one can see well-defined peaks from Chriz protein and the 5’ ends of active genes localized in them, indicating that interbands are present (Fig. 6). Six Chriz and FAIRE-seq peaks are colocalized with AQUAMARINE chromatin; between them, there lie LAZURITE chromatin fragments filled with H3K36me3 (Figs. 4, 8). In addition to these three regions, the fourth interband contains no prominent Chriz and FAIRE-seq peaks, but the 5’ end of the *CG3703* gene is located in it. Hence, we can precisely localize three gray bands with length ranging from 3.7 to 5.5 kb.

### Between bands 1E1-2 and 1E3-4

The region 1Е1-2…1E3-4 contains two black bands with a single gap between them, which can be unequivocally considered an interband; there are no gray bands in this region on the Bridges’ map (Bridges, 1938). This interband is indiscernible in cytological sections; therefore, we determine its size according to molecular maps and the positions of probes CG3021 and CG14629; they are labeled to the middle of this cytological section (Fig. 7).

The interval between the terminal ends of black bands 1E1-2 and 1E3-4 is 27.4 kb long. The four Chriz and FAIRE-seq peaks (Figs. 6, 8) are colocalized with AQUAMARINE chromatin; LAZURITE chromatin fragments filled with H3K36me3 located between them, except for one gap for which the chromatin state cannot be identified using the 4HMM because of the 6.5-kb LTR-filled break (Fig. 1F, G). Hence, we can suppose that there are two pairs of interband/gray band complexes; the bands are 2.7 and 4.7 kb long, which is below the cytological resolution threshold.

### Between bands 1E3-4 and 1F1-2

The interval between the terminal ends of black bands 1E3-4 and 1F1-2 is 28.0 kb long and contains a single gray band 1E5 on the Bridges’ map (Bridges, 1938). On the molecular map, one can see well-defined peaks from Chriz protein and the 5’ ends of active genes localized in them, indicating that interbands are present (Fig. 6). Six Chriz and FAIRE-seq peaks are colocalized with AQUAMARINE chromatin; between them, there lie LAZURITE chromatin fragments filled with H3K36me3. Therefore, we can precisely localize another gray band along with 1E5; the bands are 2.2 and 10.7 kb long.

### Between bands 1F1-2 and 2A1-2

The interval between the terminal ends of black bands is 59.2 kb long and contains two gray bands 1F3 and 1F4 on the Bridges’ map (Bridges, 1938), each corresponding to a length of 29.6 kb on average. On the molecular map, one can see well-defined peaks from Chriz protein and the 5’ ends of active genes localized in them, indicating that interbands are present (Fig. 6). Eight Chriz and FAIRE-seq peaks are colocalized with AQUAMARINE chromatin; between them, there lie LAZURITE chromatin fragments filled with H3K36me3, except for one gap. For this region, the chromatin states cannot be identified using the 4HMM because of the gap comprising the 4.7 kb LINE repeat. Hence, we can precisely localize seven “gray” bands, with lengths ranging from 2.7 to 10.5 kb, on the molecular map.

## Discussion

Interphase and mitotic chromosomes have a complex system of DNA and chromatin packing (Mortin and. Sedat, 1982; Mathog et al., et al., 1984; Sedat et al., 2022a, b; McDonald et al., 2024). As a result of this, the well-known transverse banding pattern in *Drosophila* is formed in interphase polytene chromosomes. Such striping has been found in human interphase chromosomes (Sedat et al., 2022b). Therefore, the striping could be considered as universal element of genome organization at least in *Drosophila* and humans.

Depending on the degree of compactness of chromatin packing in different chromosomal structures, two types of bands (black and gray) and interbands are distinguished. Black bands are characterized by the densest DNA packing (160-220 times), gray bands have an intermediate density (25-35 times) and interbands are maximally decondensed (5-10 times) (Kozlova et al., 1994; Vatolina et al., 2011). Enormous differences were found in the genetic organization of these three structures: the inter bands contain origin recognition complexes that initiate replication and promoters of housekeeping genes, the bodies of which are located in neighboring gray bands. The black bands contain clusters of developmental genes. The developmental and housekeeping genes differ in many genetic features: nucleotide motifs in promoters, sizes of introns, exons, intergenic spacers, and the number of transcription start points (Zhimulev et al., 2024).

In this study, we have used the numerous state-of-the-art data on chromatin structure, gene localization, and methods for quantifying their activity. We managed to plot a detailed map of the region 1AF in interphase polytene chromosomes as well as developmental and housekeeping genes. We have determined the lengths and genetic content of the bands containing the developmental genes and regions or individual genes localized in gray bands or groups thereof, where more precise mapping is infeasible. The lengths of cytological structures and genes localized in them are presented in Table S1.

A comprehensive method for combining the physical map of the *Drosophila* genome and the cytological map of the interphase polytene chromosome has been elaborated in this study. When performing molecular and cytological mapping, we utilized the data on Chip-seq distribution and immunofluorescence localization of interband-specific Chriz protein simultaneously with open chromatin localization using such techniques as FAIRE-seq, ChIP-seq of distribution of antibodies specific to H3K36me3 histone modification, a marker of transcription elongation, and the 4HMM of four chromatin states that allowed detection of open and closed chromatin.

The map of mutual arrangement of genes, genomic elements, and cytological structures with respect to the long region 1AF of polytene chromosome has been created (Fig. 9). Figure 9 also shows the map of chromosome structures currently available in databases (FlyBase) compared to the findings obtained in this study.

Comparison of molecular maps demonstrates that band boundaries coincide in none of the cases, although sometimes they overlap.

**Figure 9.**
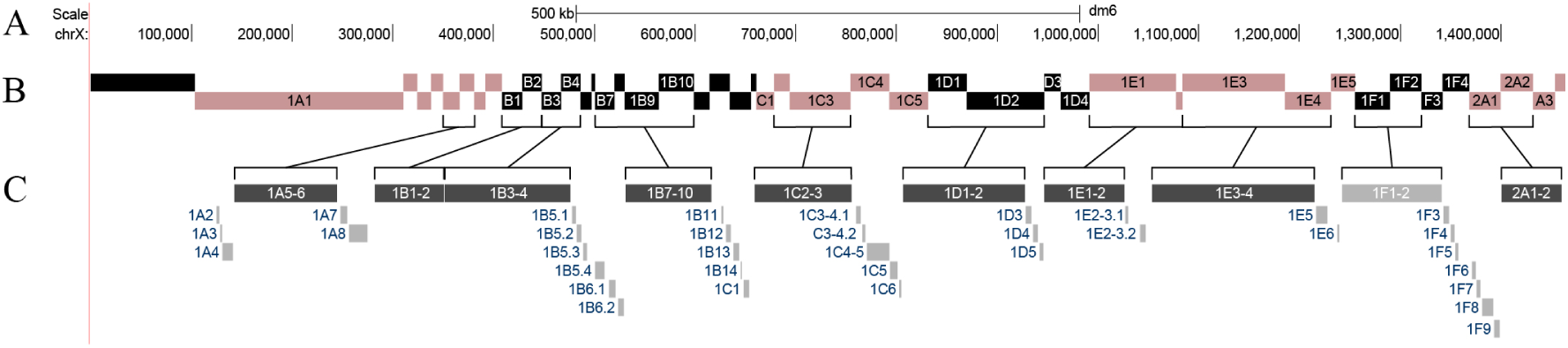
Comparison of the positions of bands and interbands available in the UCSC dm6 database and those mapped in this study. (A) Scale and molecular coordinates, UCSC release dm6. (B) Band positions in accordance with the FlyBase data (D. melanogaster version 6.02). (C) Schematic arrangement of bands and interbands in accordance with the data reported in the present paper.

Having compared two molecular cytogenetic maps, one can infer that:

1. Band sizes on the two maps differ drastically: on the FlyBase map (Fig. 9B), almost all the bands have the same length and do not demonstrate DNA lengths corresponding to certain bands in terms of their cytological pattern, whereas the banding pattern observed in this study reveals matching of band size and correlates with their length (the amount of DNA in them).
2. On the FlyBase map, there is no subdivision into black (positions of developmental genes) and gray (housekeeping genes) bands. Furthermore, differences in their lengths are not taken into account. DNA content in gray bands is manifold lower than that in black bands (see Fig. 8G, H).
3. On the FlyBase map, there are neither positions nor mentions of the critical genomic elements: interbands where promoters of the 6562 genes of housekeeping genes and protein Origin Recognition Complexes (ORCs) are located (Fig. 8) (Zhimulev et al., 2024). In other words, the most significant portion of the genome is not represented in the “checkerboard” pattern.

The results of this study focusing on the genome organization in the region 1AF are summarized in two tables (Table 2, Table S1). Table S1 lists the detailed mapping data for the region 1AF. The length and precise molecular coordinates determined using the 4HMM are specified for each interband, gray and black band (inaccuracy ≤ 200 bp). The degree of overlapping with ORC2 in salivary gland and S2 cells are additionally provided for each structure, and the genes residing in this interval are listed (the number of genes was calculated if any portion of the gene was present in the structure, regardless of whether it was coding or non-coding; 5’ and 3’ ends and introns were given the same value, regardless of the overlapping percentage). A total of eight black bands, a large light-colored band (1F1-2), and one black band (2A1-2) confining the region at the proximal side were detected in the region under study (a total of ten bands were included in the sample). In the gaps between the large bands, there were series of thin gray bands alternating with interbands. The region contains a total of nine such series, with 35 gray bands detected in them. Hence, the series contain complexes following each other, each of them comprising at least one gene having its 5’ end in the interband, whereas the gray band is formed by the gene body.

The genetic composition of chromosome structures in the 1AF region has been studied thoroughly (Table 2).

**Table 2.** Gene size and distribution within the interphase chromosome structures according to physical mapping data.

| Region 1AF | Number of structures | Total length of a single structure, kb | Average length of a single structure, kb | Total number of genes in the structure | Average number of genes per structure | Average number of genes per 1 kb |
| --- | --- | --- | --- | --- | --- | --- |
| telomere | 1 | 131.21 | 131.21 | - | - | - |
| interbands | 45 | 125.00 | 2.78 | 87 | 1.93 | 0.70 |
| gray bands | 35 | 197.57 | 5.64 | 62 | 1.77 | 0.31 |
| black bands | 10 | 795.65 | 90.37 | 65 | 6.50 | 0.08 |
| gaps | 2 | 10.20 | 5.10 | 1 | 0.50 | 0.10 |

The telomere is 131.2 kb long and is an exception, since it does not contain annotated genes. Due to the coordinates precisely determined using the 4HMM, it turned out that on average, the interbands within the region under study are 2.78 kb long; gray bands, 5.64 kb long; and black bands, 90.37 kb long, which is consistent with the experimental data obtained earlier (Beermann, 1972; Kozlova et al., 1994; Vatolina et al., 2011). Hence, it turned out that most genes reside in large black bands (6.50), the average number of genes per kb being minimal (0.07). An almost identical number of genes are on average located within interbands and gray bands (1.93 and 1.77, respectively); however, the gene density per kb is higher than that in gray bands (0.70 vs 0.31) (Table 2). This estimate is consistent with the data that the developmental genes are long and are encompassed by black bands, whereas the interbands contain the maximum number of active but short housekeeping genes.

Being 131.2 kb long, the telomere is visualized by optical microscopy as a light-colored loose structure, whereas black bands of a similar size are dense, large, and well-discernible. This may indicate that DNA or nucleosome repeat copies are under-represented within the telomere.

The data on genetic organization of both types of bands located in these structures were also obtained; it was calculated according to the presence of any gene in the structure, regardless of whether it was coding or non-coding (introns, as well as the 5’ and 3’ ends were given the same value, regardless of the percentage of overlapping with the chromosome structure under consideration).

Hence, it turned out that the largest number of genes reside in black bands (6.50 per band), whereas the average number of genes per kb is the smallest. It implies that the number of genes in black bands is small, and the genes are rather long.

An opposite situation is observed for the interband/gray band complexes: each of them contains almost two genes (1.93), and their number per kb is 0.70, meaning that the genes are short. The number of genes in gray bands is comparable to that in interbands (1.77), but the number of genes per kb is twice lower: 0.31 (Table S1).

Therefore, this study allowed us to present, in a noncontradictory form, all the data that can be presented for the arrangement of the interphase (i.e., functioning) genome: the housekeeping and developmental genes, their sequence and positions related to length of cytological structures (bands and interbands), as well as such genomic elements as ORC2, interband-specific protein Chriz, and H3K36me3 histone modification. Since it is important that the chromosome structure and gene activity are compared across the same cell type, most data were obtained for salivary glands.

The boundaries of chromosome structures and genes encompassed by them were identified very precisely for the first time to complete the discussion on whether one band contains a single gene. The findings provide good information about the two types of chromosomes: in the interphase dividing cells as well as in polytene chromosomes and salivary glands (Vatolina et al., 2011). The boundaries of the bands that can be visualized by FISH strictly correspond to the boundaries of 4HMM chromatin states comprised by these structures. Localization of chromatin structures and states corresponding to them is strictly consistent with the results of a previous study.

Research based on 3C, Hi-C, and micro-C high-throughput sequencing methods for studying chromatin conformation has become commonly performed in the 2000s (Lieberman-Aiden et al., 2009). These methods are based on formalin fixation of contacts between chromatin regions existing in the cell nucleus; chromatin is then solubilized and fragmented; the interacting DNA regions are ligated together to create the genomic library of chimeric molecules. Once fragments contacting each other more frequently were selected, topologically associated domains (TADs) can be formed. The relative abundance of chimeric fragments and ligation products correlates with the probability that the respective chromatin fragments interact in the 3D space across the entire cell population. Whereas the 3C method focuses on analyzing the set of predetermined genomic loci, the Hi-C technique enables profiling of the “all-versus-all” interaction by labeling the entire fragmented chromatin with a biotinylated nucleotide prior to ligation.

R. White (2012) hypothesized that the domains are linked to the bands in polytene chromosomes. In our study, we demonstrated for two dense bands 10A1-2 and 10В1-2 that TAD loops relatively corresponded to these black bands. We have run through polytene chromosome ∼ 1100 regions containing RUBY chromatin fragments and revealed good matching between black bands and TADs (Zhimulev et al., 2014).

Quite a few studies have focused on TADs, which have been drawing close attention of scientists specializing in many disciplines (Eagen et al., 2015; Ulianov et al., 2016; Stadler et al., 2017; Ball et al., 2023; Bing et al., 2024; Fishman et al., 2025). Therefore, we decided to verify how well TADs match the structures of the interphase polytene chromosomes within the sufficiently long region that has been studied well.

Figure 10 shows the results of analyzing the DNA contact points upon identification and isolation of TADs only in three papers. The upper panel of Fig. 10 shows heatmaps of chromatin contact frequencies generated from the processed primary Hi-C data during TAD formation (see Methods section). The middle panel presents insulation score profiles, which quantify the degree of local insulation between adjacent chromatin regions and are commonly used to infer TAD boundaries. The lower panel displays the corresponding cytological map, indicating the positions of black and grey bands along the chromosome.

**Figure 10.**
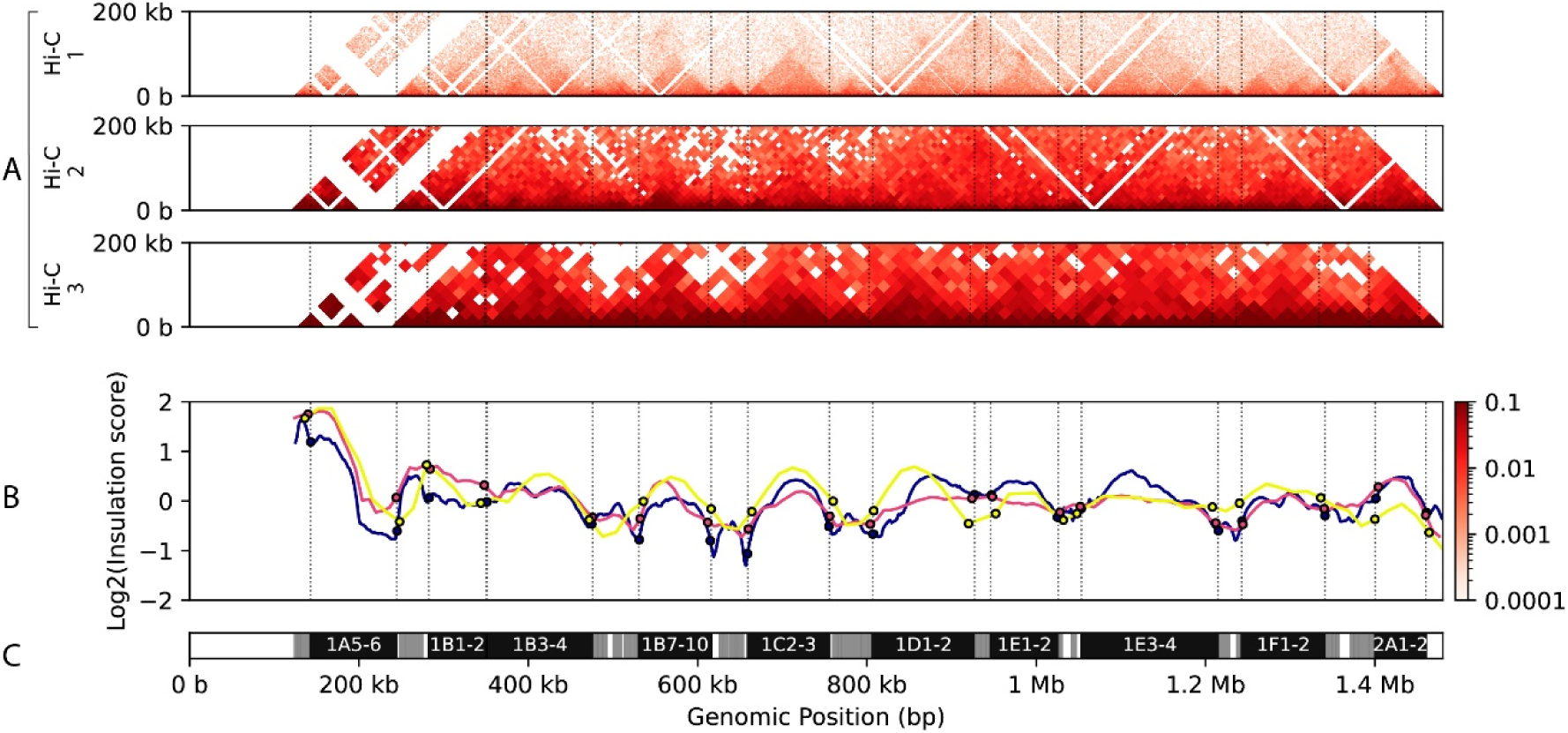
Comparative Hi-C contact maps, insulation score profiles, and cytological banding in the 1AF region. (A) Heatmaps of normalized Hi-C contact frequencies for three datasets: 1 – (Bing et al., 2024) (GSE263229) at 2 kb resolution (top), 2 – (Stadler et al., 2017) (GSE100370) at 8 kb resolution (middle), and 3 – (Eagen, 2015) (GSE72512) at 16 kb resolution (bottom). Vertical dashed lines delimit the boundaries of cytologically defined black bands. (B) Log₂ transformed insulation score profiles computed with sliding windows of 40 kb for (Bing et al., 2024) (blue), and 80 kb for both (Stadler et al., 2017) (red) and (Eagen et al., 2015) (yellow). Black circles mark the points where each insulation profile intersects a black band boundary. (C) Cytological map of the 1AF region showing positions of black (dark) and grey (light) bands along the chromosome.

Stadler et al. (2017) pointed to the lack of high-resolution data correlating the structural features of polytene chromosomes with specific genomic coordinates. In this study, we used novel approaches to construct a detailed map of interbands and two types of bands (black and gray) in the region 1AF. Comparison of the Hi-C data from multiple publications and cytological maps (Fig. 10) with our newly generated map of the region 1AF revealed that the boundaries between black and gray bands are well-discernible in the map of TADs and are characterized by local minima of the insulation score, the hallmark feature of TAD boundaries. Hence, two TAD categories can be differentiated for the utilized resolution of the Hi-C maps: those corresponding to black bands and groups of gray bands. However, all the groups of faint gray bands and interbands are typically represented by a single TAD loop. It is fair to assume that higher resolution of the Hi-C data (< 1 kb) will make it possible to single out individual gray bands rather than their groups.

In conclusion, genome-wide mapping of the Hi-C data, molecular biological identification of the chromatin structure, and cytological mapping are well-matched, especially for the genomic region harboring developmental genes localized within black bands.

## Article and author information

### Author details

**Varvara A Khoroshko**

Institute of Molecular and Cellular Biology, Siberian Branch of the Russian Academy of Sciences, Novosibirsk, Russia

**Anton V Tsukanov**

Federal Research Center Institute of Cytology and Genetics, Siberian Branch of the Russian Academy of Sciences, Novosibirsk, Russia

**Nadezhda E Vorobyeva**

Institute of Gene Biology, Russian Academy of Sciences, Moscow, Russia

**Tatyana Yu Vatolina**

Institute of Molecular and Cellular Biology, Siberian Branch of the Russian Academy of Sciences, Novosibirsk, Russia

**Viktor G Levitsky**

Federal Research Center Institute of Cytology and Genetics, Siberian Branch of the Russian Academy of Sciences, Novosibirsk, Russia

**Igor F Zhimulev**

Institute of Molecular and Cellular Biology, Siberian Branch of the Russian Academy of Sciences, Novosibirsk, Russia

## Author Notes

Competing interests: No competing interests declared

## Acknowledgments

The authors are grateful to Tatyana D. Kolesnikova for providing photographs of some regions of the X chromosome.

---

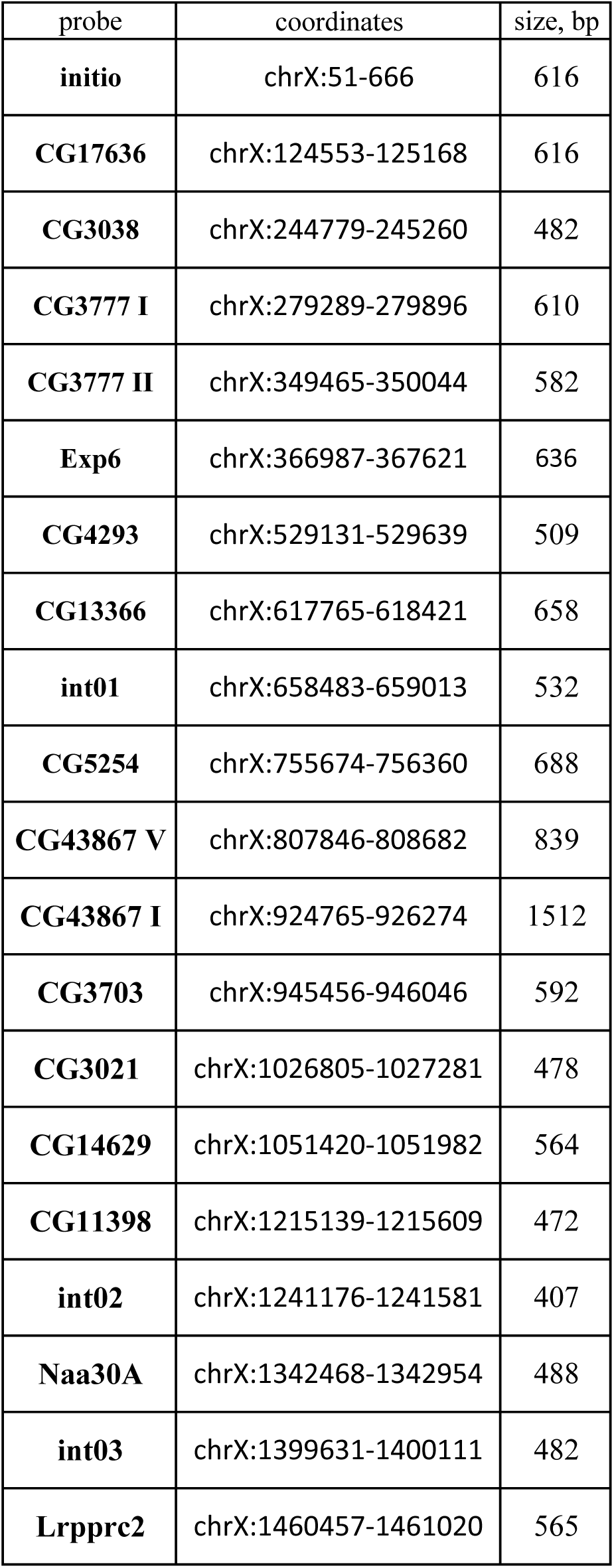
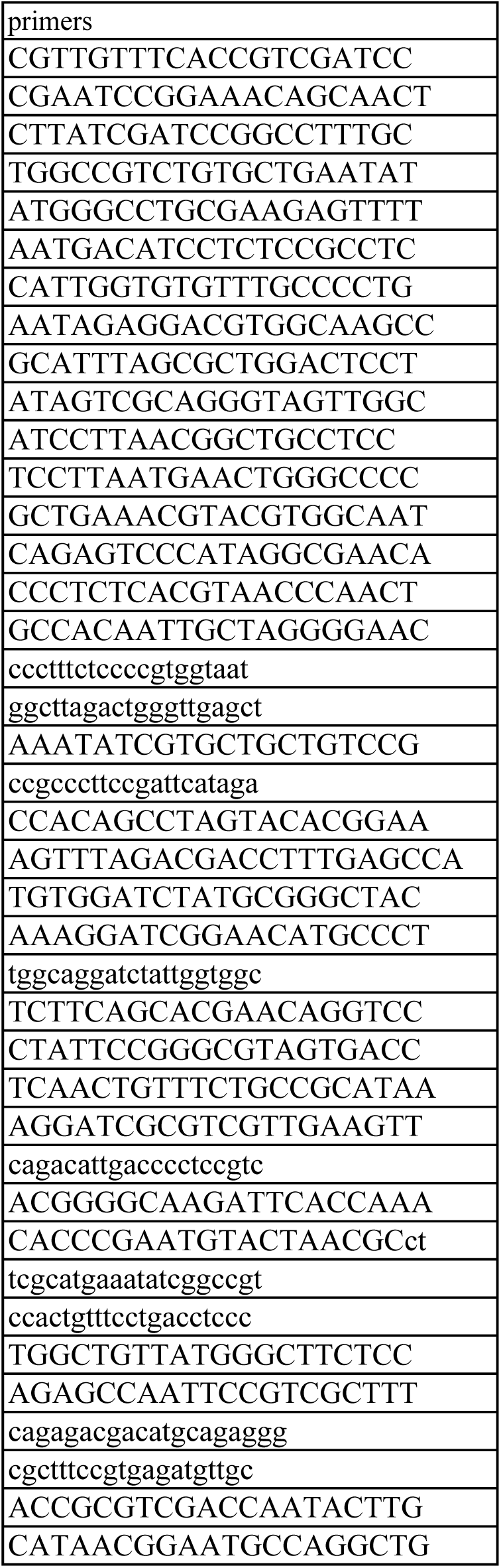

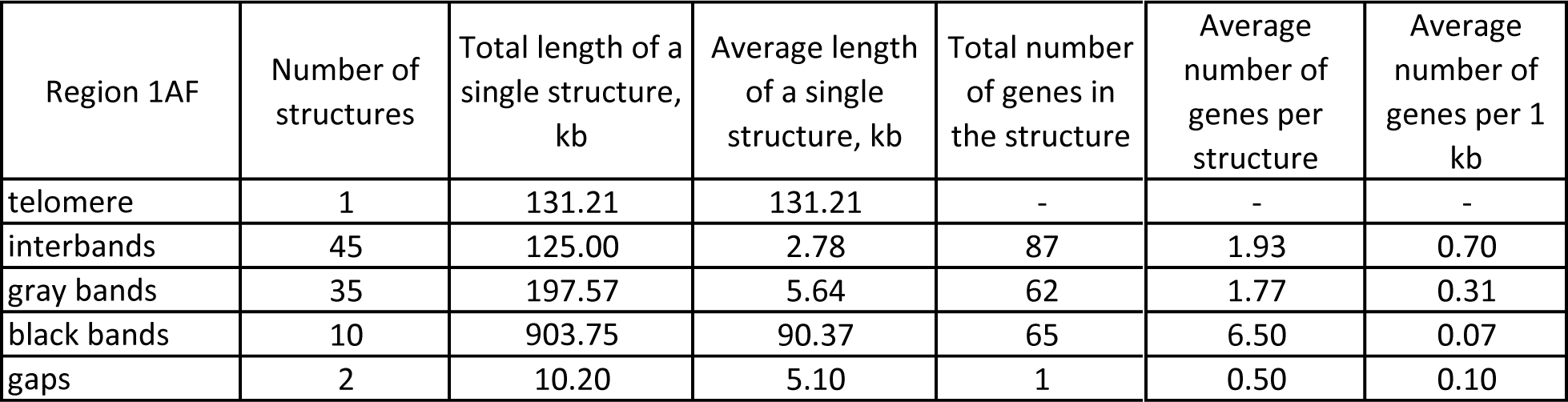

